# IL6 mediated cFLIP downregulation increases the migratory and invasive potential of triple negative breast cancer cell

**DOI:** 10.1101/2024.01.14.575471

**Authors:** Samraj Sinha, Rajdeep Roy, Nilesh Barman, Purandar Sarkar, Abhik Saha, Nabendu Biswas

## Abstract

c-FLIP (cellular FLICE-Like Inhibitor of Apoptotic protein) which belongs to the negative regulators of the death receptor pathway, control apoptosis in a number of cancers. literatures suggest that its overexpression facilitate cancer progression and also helps in the drug resistance. However, there is a very few data available to show how it control metastasis. Moreover, how cFLIP is regulated during the progression of cancer into advanced stage and its mechanism of action on tumor cell migration and invasion is yet to be known. Our TCGA data analysis has shown that cFLIP is downregulated in many cancers, including breast cancer, especially at the later stages. when, we analysed and compared early-stage breast carcinoma cell line MCF7 (luminal) and advanced stage triple negative cell line MDAMB-231, which is highly metastatic, we found a negative correlation between cFLIP expression and metastasis. Our results indicated that Il6, one of the most prominent cytokines inside tumor microenvironment, activated p38 and the later helped in cFLIP downregulation in MDA-MB-231 cell line. Moreover, we have found that cFLIP negatively regulated autophagy and this autophagy downregulation resulted in decrease in metastasis. Thus, we have shown, for the first time, a complete interconnecting pathway in which IL6 mediated p38 activation directly influences metastasis by regulating autophagy via cFLIP downregulation.

## Introduction

c-FLIP (cellular FLICE-Like Inhibitor of Apoptotic protein), the structural homologue of procaspase-8, binds to the Fas-associated death domain (FADD) at the death-inducing signalling complex (DISC) and thus inhibits FasL-mediated apoptosis (1,2). Antagonising cFLIP’s anti-apoptotic activity is considered as a potential strategy to treat cancer cells (3). cFLIP has been identified to be overexpressed in a significant number of patients in non-Hodgkin’s lymphoma (4), bladder urothelial carcinomas (5), Colon carcinoma (6). Recently we have shown that cFLIP plays a pivotal role in the apoptosis of imatinib-sensitive and imatinib-resistant CML cells and it is differentially regulated by ROS in drug sensitive and resistant cells (7,8). However, there is very limited knowledge of how cFLIP down regulation plays a role in the progression of several cancers, including breast cancer.

Triple-negative breast cancer (TNBC), which lacks expression of oestrogen receptor (ER), progesterone receptor (PR) and human epidermal growth factor receptor 2 (Her-2), has a poor prognosis and is the most aggressive subtype of breast cancer. Metastasis is one of the major hindrances for the treatment of TNBC as it is mostly detected at advanced stages. Spreading of breast cancer cells has been mainly found in the lymph nodes, bones, lungs, liver, and brain (9). Therefore, targeting metastasis is an ever-growing area of research in TNBC.

Several factors play a role in initiating or regulating this metastasis with progression of tumour. Recent studies suggested that cytokines play important role in the metastasis (10,11) as cytokine profiles changes inside tumour microenvironment with the progression of tumour. Among these cytokines, IL6 represents as one of the most important cytokines, which has a pivotal role in normal homeostasis and disease augmentation (12). IL-6 is produced in the tumour by infiltrating immune cells, by the tumour cells themselves, and by stromal cells. Thus, tumour-associated macrophages, granulocytes, and fibroblasts, as well as cancer cells, are all primary sources of IL-6 (13,14). Canonically IL6 is known to bind to a dimer of IL6-R and GP130 complex and activate the JAK/STAT pathway (15). IL6 plays a crucial role in cancer progression by blocking immune cells from recognising the cancer cells. A number of studies demonstrated that IL6 imparts chemoresistance against drugs like tamoxifen, doxorubicin in HER2+ breast cancer cell lines. Other evidences also report its’ pro-tumorigenic effect on TNBC lines (16).

Here, in this project, we investigated the roles of cFLIP downregulation in the metastasis of breast cancer. Our result concluded that cFLIP is downregulated in the advanced stage of breast cancer and this downregulation helps in metastasis. IL6 mediated p38 activation played a key role in cFLIP downregulation. Moreover, we have also revealed that upregulation of autophagy as a result of downregulation of cFLIP is involved in the increase in migration and invasion potential of triple negative breast cancer cells.

## MATERIALS

Antibodies against proteins c-FLIP, p-ERK, p-JNK, p-P38, ITCH, p62, LC3B, ATG12, Vimentin, E-cad, ZEB-1, Actin, snail, twist, MMP9, p-cJun anti-Rabbit IgG & anti-mouse HRP-linked antibodies, Control siRNA, p38 siRNA were bought from Cell Signaling Technology (Denver, Massachusetts, USA). Transwell insert plates were taken from ThermoFisher Scientific and matrigel from Corning. Fetal Bovine Serum (FBS) Standard (origin South America) was acured from Gibco.Life Technologies, USA. iScript Reverse Transcription Supermix and SsoFast Evagreen Supermix were got from BIO RAD (USA) for performing RT-qPCR. Recombinant IL6 protein was purchased from BIOLEGENDS (570802).

## METHODS

### Cell Line and cell culture

MDA-MB-231, MDA-MB-468, MDA-MB-453, MCF7 cell lines were purchased from National Centre for Cell Science, India. The cells were maintained in DMEM medium to which 1% penicillin-streptomycin and 0.1% gentamycin and 10% FBS was added. All cells were cultured at 37^0^C humidified condition with continuous supply of 5% carbon dioxide (CO_2_).

### Western blot analysis

After the cells were treated as mentioned in the respective figure legends, 1X PBS was used to twice wash the cells before lysing them in 1X RIPA (CST, Cat. no 9806S), containing protease inhibitor cocktail. Cells were vortexed and sonicated for 15 seconds by keeping them alternatively on ice and then cell lysate was isolated using high speed centrifuge at 15000 rpm for 15 minutes. The protein concentration was measured using by Lowry (Folin-Ciocalteau) method as standard protocol. Then the lysates were loaded to SDS-PAGE after boiling them in Laemmli buffer at 95^0^C for 5 minutes. After the electrophoresis the gel was blotted to PVDF membrane. 5% non-fat dried milk powder in TBST were used to block the membrane for around 2 hours and incubated in respective antibodies overnight. For detection of the primary antibody, the membrane was incubated in HRP linked corresponding secondary antibodies diluted in TBST for one hour. Blot development was done using BIO RAD Clarity Western ECL substrate. BIO RAD ChemiDoc MP Imaging System and by ImageLab 5.0 software images were obtained.

### Gene overexpression

Ectopic expression of cFLIP was done with pcDNA3-cFLIP (a kind gift from Dr. Mikhiko Naito, National Institute of Health Sciences, Tokyo, Japan) and the empty pcDNA3 was used as control. Transfection was done using Lipofectamine 3000 as mentioned in the manufacturer’s protocol. After 48 hours or else as indicated cell lysate if not otherwise mentioned, was isolated and subjected to Western blot to confirm the overexpression and its further downstream target expressions.

### Gene knock down assay

About 2 million cells were seeded in each well of six-well plates and kept overnight in serum and antibiotics free optimal media. Following day, respective siRNA oligo transfection was performed using Lipofectamine 3000 (Invitrogen, Carlsbad, California, USA). After 48 hours of transfection if not otherwise mentioned, western blot was performed with the isolated lysates to confirm the knockdown of the candidate gene.

### Real time PCR

TRIZOL reagent was used to isolate the RNA from the indicated cells. By agarose gel electrophoresis and Synergy H1 Microplate Reader were used to check the quality and quantity of RNA respectively. iScript Reverse Transcription Supermix was used to prepare cDNA from isolated total RNA. Now Real time PCR was performed by using SsoFast Evagreen Supermix (BIO-RAD) and analysed in CFX96 Touch Real-Time PCR Detection System (BIO-RAD) Minimum 40 cycles were ran for PCR for reaching total saturation and the Cq values were considered for further analysis. Actin was used as housekeeping gene to normalize the mRNA levels and the data were presented graphically as expression ratio to Actin.

### Immunofluorescences microscopy

Round glass bottom dish (Invitro Sci.) was used for seeding around 0.2 million cells. Cells were fixed using 10 % PFA for 15-20 minutes. BSA containing NP-40 buffer was used to wash the cells for about two hours and then the primary antibodies (1:25) was added and incubated overnight. Next day cells were again washed using NP-40 buffer for 2 hours and incubated with DAPI and secondary antibody (Alexa Fluro 594, CST) in 1:400 ratio diluted in NP-40 buffer. Again, cells were washed by the NP-40 wash buffer for around two hours. All images were obtained using Leica DMi8 fluorescence inverted microscope.

#### Cell scratch test

Cells were seeded in 6-well plate or/and experiments were performed as indicated, once it reached good confluency, a cross shaped scratch was done using a sterile 10 μl pipette tip. These cells were kept in FBS-free culture medium (to nullify the effect of growth factors on cell migration). Time dependant images as indicated in figures of the scratches were captured in LAS EZ version 3.4 microscope. The width of different positions of the scratch was assesed with the distance measurement tool of ImageJ and represented in the form of bar graphs.

#### Transwell migration assay

Transwell migration assay was done using transwell inserts of 8 μm pore diameter. Equal number of cells were seeded in serum-free medium in upper chamber (in case of invasion assay the wells were covered with matrigel and left overnight before cell seeding) and treatment was done as mentioned and media with FBS was added to the lower chamber. After indicated period of incubation, the cell fixation was done by methanol and stained by Giemsa. Then the cells of the lower surface were examined with LAS EZ version 3.4 microscope. Photos of four random fields were taken for using them in counting purpose and the average number of cells were used to measure the migration capacity.

### Statistical Analysis

Statistical significance was determined by the two-tailed paired Student’s t-test in all experiments in this study. The data are presented as means ± standard deviation (SD). Values of p<0.05 were considered statistically significant. Asterisks indicate the statistical significance as follows: * p<0.05. All the western blot analysis data are representative blots of three independent experiments.

## RESULTS

### cFLIP expression is downregulated in metastatic cancers with respect to normal tissues

Previously we have shown that cFLIP levels remain higher in drug resistant-CML cells as compared to the drug-sensitive counterpart (unpublished data). Here, we aimed to evaluate cFLIP expression profile in other cancers with a special emphasis on breast cancer. The RNA-seq analysis of TCGA database revealed that cFLIP expression is significantly downregulated in different cancers including breast carcinoma when compared to the normal tissues (Figs. 1A and 1B). Since the TCGA data showed the downregulation of cFLIP in cancer samples as compared to the normal tissues, we extended our analysis into the early and late stage cancer to check its temporal expression pattern. The analysis demonstrated that cFLIP expression is strikingly diminished in metastatic stage with respect to non-metastatic tumour and normal tissues (Figs. 1C and 1D). We therefore hypothesized that lower levels of cFLIP may be connected with poor clinical results owing to aggressive metastatic conditions. Indeed, analysis of data derived from luminal breast cancer as well as TNBC patient samples revealed, lower levels of cFLIP (FLAME) expression being correlated with poor patient outcomes as measured by diminished overall survival (OS) and disease-free survival (DFS) (Fig. 1E).

**Figure 1:**
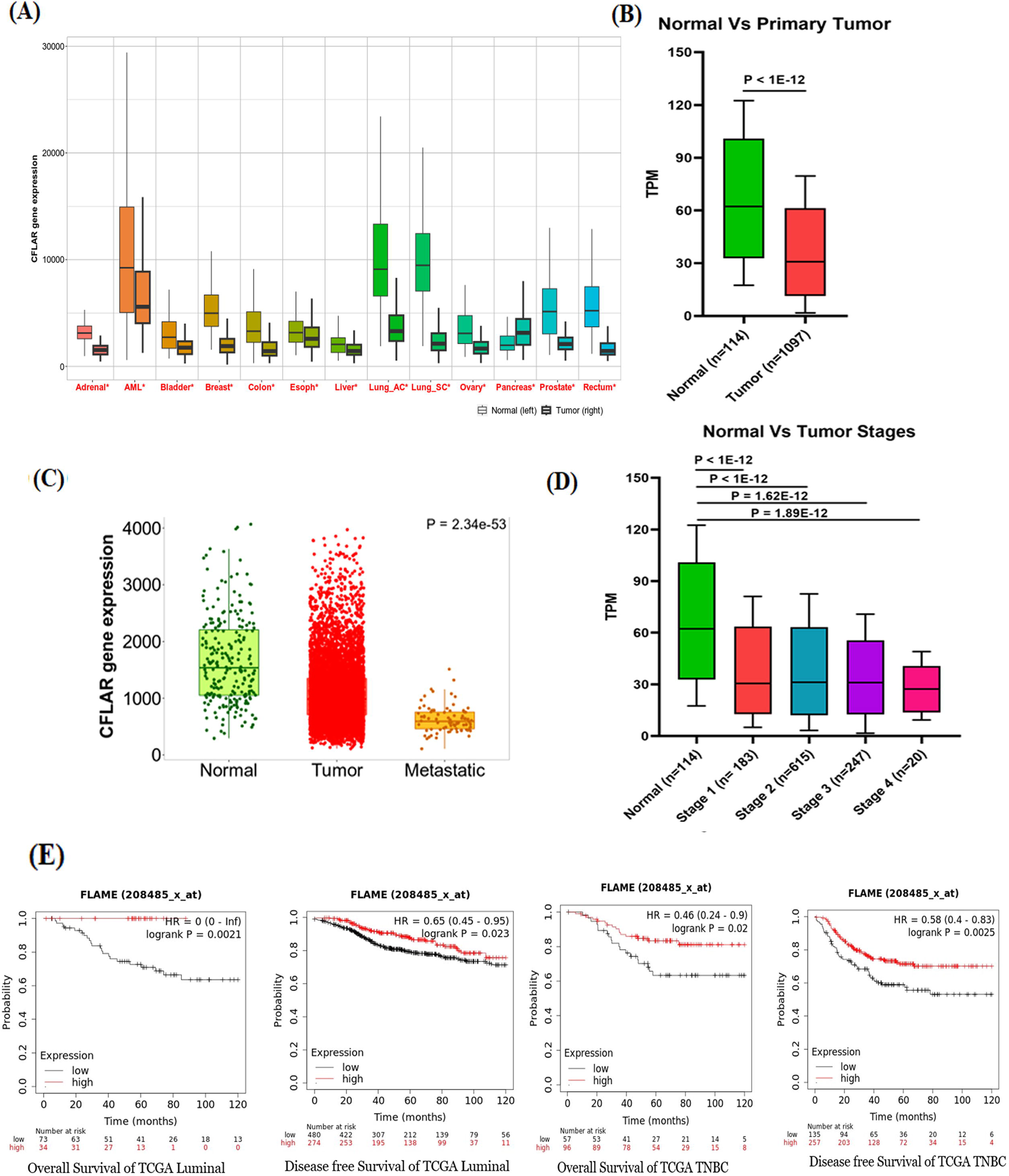
cFLIP expression is downregulated in metastatic breast cancers with respect to normal tissues. (A) cFLIP expression profile in different cancer vs normal tissues. (B) cFLIP expression levels in cancer vs normal breast tissues. (C) cFLIP expression levels in normal vs cancer vs metastatic breast tissues. (D) cFLIP expression levels in normal vs cancer stages of breast tissues. (E) Kaplan-Mayer’s plot showing survival probability in luminal and TNBC breast cancer types.

### Correlation of the migratory ability of different breast cancer cell lines with cFLIP expression

It is well known that the metastatic ability of the cancer cells causes the aggressive spread of the tumour in patients and declines the survival ability. TCGA analyses prompted us to further extend our study into cell line model, where we utilized four breast cancer lines – including luminal type MCF7 along with three TNBC lines MDA-MB-231, MDA-MB-468 and MDA-MB-453, which were derived from more advanced metastatic stage of breast cancer patients. In order to assess the metastatic ability of these cell lines, we conducted scratch assay, trans-well migration and 3D Matrigel invasion assays. (Fig. 2A, 2B and 2C, respectively). In scratch assay, 24h post-treatment the wound area as expected was significantly reduced in MDA-MB-231and MDA-MB-468 lines as compared to MCF7 (Fig. 2A). However, to our surprise, wound area was not noticeably reduced in case of the third member of aggressive TNBC cell line MDA-MB-453.

**Figure 2:**
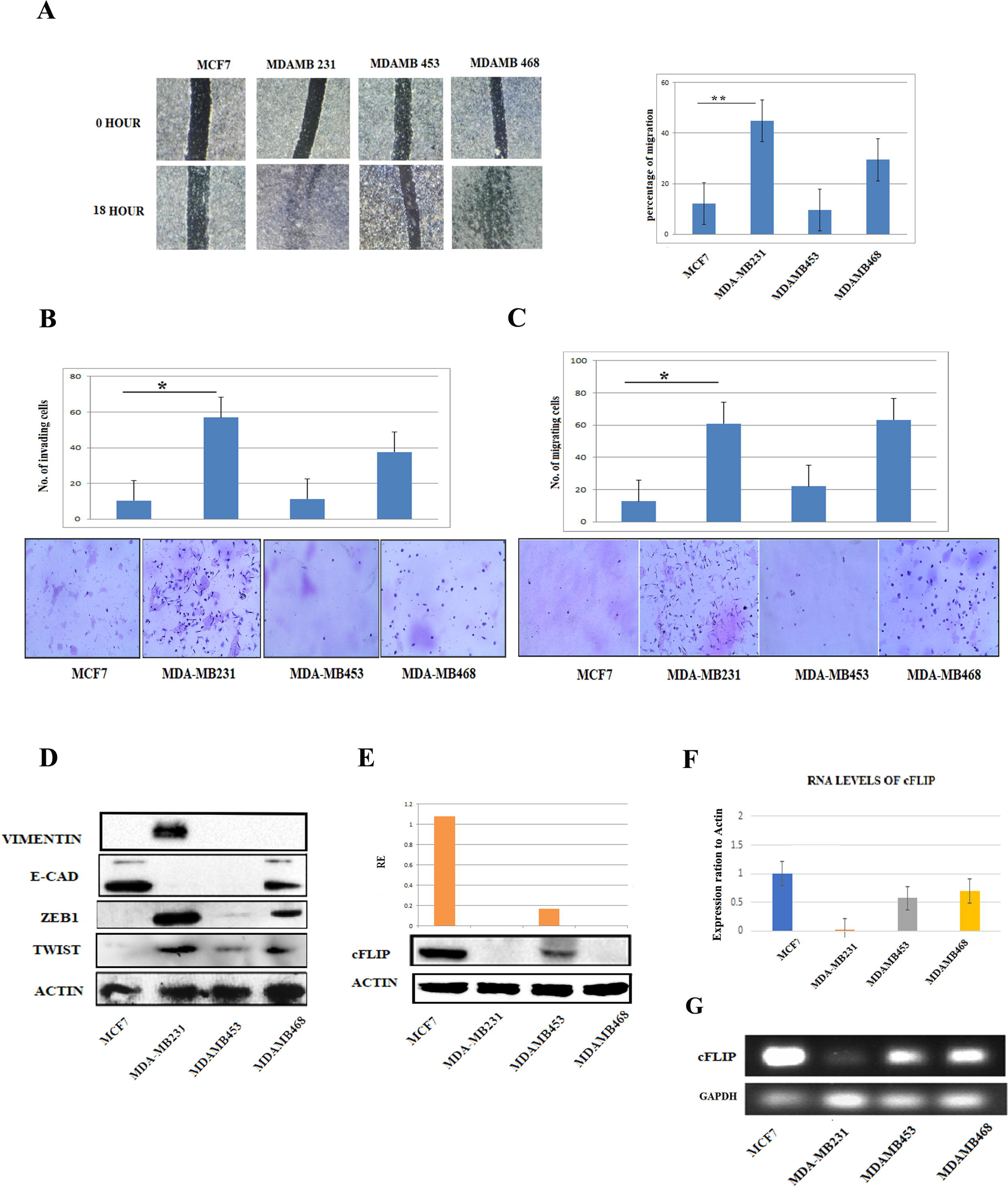
Correlation of the migratory ability of different breast cancer cell lineas with cFLIP expression in them. (A) Scratch assay was performed according to the protocol for MCF-7, MDAMB-231, MDAMB-453, MDAMB-468. Representative photographs of one random field is provided and the wound area was represented in form of bar graph with mean ± SD. (B) Invasion assay was performed in the standard protocol with the mentioned cell lines and four random representative fields were photographed. Cells were counted and represented in form of bar graphs with mean ± SD. (C) Transwell migration assay was performed in the standard protocol with the mentioned cell lines and four random representative fields were photographed. Cells were counted and represented in form of bar graphs with mean ± SD. (D) The expression of epithelial to mesenchymal transition markers were analysed using western blot from the four mentioned cell lines. For loading control γ Actin was checked. Densitometric analysis data was represented graphically normalised with actin and indicated ± SD. (E) The protein levels of FLIP was measured using western blot analysis from four mentioned cell lines. For loading control γ Actin was used. Densitometric data were presented graphically normalised to actin. (F) Relative levels of endogenous FLIP mRNA in the indicated cell lines were evaluated by Real time PCR and graphically denoted as fold change with respect to actin. Data represent mean ± SD. Expression levels of cFLIP mRNA was checked with the help of semi-quantitative PCR with respect to GAPDH mRNA was represented.

Next, we evaluated the invasion pattern of each cell line using invasion assay. The results demonstrated that MDA-MB231 cell number was most in the lower chamber, whereas other TNBC lines MDA-MB-468 showed a good number of cells in the lower chamber (Figs. 2B and 2C). However, the luminal type MCF7 cells and TNBC cell line MDA-MB-453 were found significantly lower in number. These data indicated that MDA-MB-231 cells show the most metastatic ability and MCF7 the least. In order to confirm the results, the expression of the epithelial to mesenchymal transition (EMT) markers were checked by western blot analysis. The results reflected that the mesenchymal markers Vimentin, Zeb-1 and Twist increased most in MDA-MB-231 line, whereas the epithelial marker E-cad is significantly elevated in MCF7 line (Fig. 2D). Overall, the EMT markers further indicated that MDA-MB-231 line is most aggressive and MCF7 the least.

Next, we wanted to correlate cFLIP expression with the metastatic potential of these lines. Both western blot and qPCR analysis showed that cFLIP expressions are negligible in TNBC lines including MDA-MB-231 and MDA-MB-468, while it remains high in luminal line MCF7 (Figs. 2E and 2F). Interestingly, despite being advanced stage cell, MDA-MB-453 line showed moderate expression of cFLIP. However, the migration and invasion assays impeccably corroborated our hypothesis correlating lower cFLIP expression with higher metastatic potential.

### cFLIP exerts a negative impact in migration and invasion properties of breast cancer cells

Given the inverse correlation of cFLIP expression with metastatic ability, cFLIP was knocked down using cFLIP-specific siRNA in MCF7 cells and scratch assay and trans-well migration assays were performed using scramble siRNA as control. In scratch assay, cFLIP knockdown significantly elevated (∼2 fold) MCF7 cell migration at 26 h as compared to the scramble siRNA-transfected MCF7 cells (Fig. 3A). Similarly, cFLIP knockdown cells showed approximately 2 fold increase in individual cell motility in matrix-specific transwell migration assay (Fig. 3B). To confirm the cFLIP knockdown, a part of the control and cFLIP siRNA transfected MCF7 cells from above-mentioned experiments were subjected to western blot analysis, which showed a marked knockdown at the protein level (Fig. 3B). In agreement with the increased cell migration/invasion ability in cFLIP knockdown MCF7 cells, western blot analysis of EMT markers further demonstrated decreased epithelial marker E-cad and increased mesenchymal markers Snail and Twist expressions in cFLIP knockdown MCF7 cells as compared to the control line (Fig. 3C).

**Figure 3:**
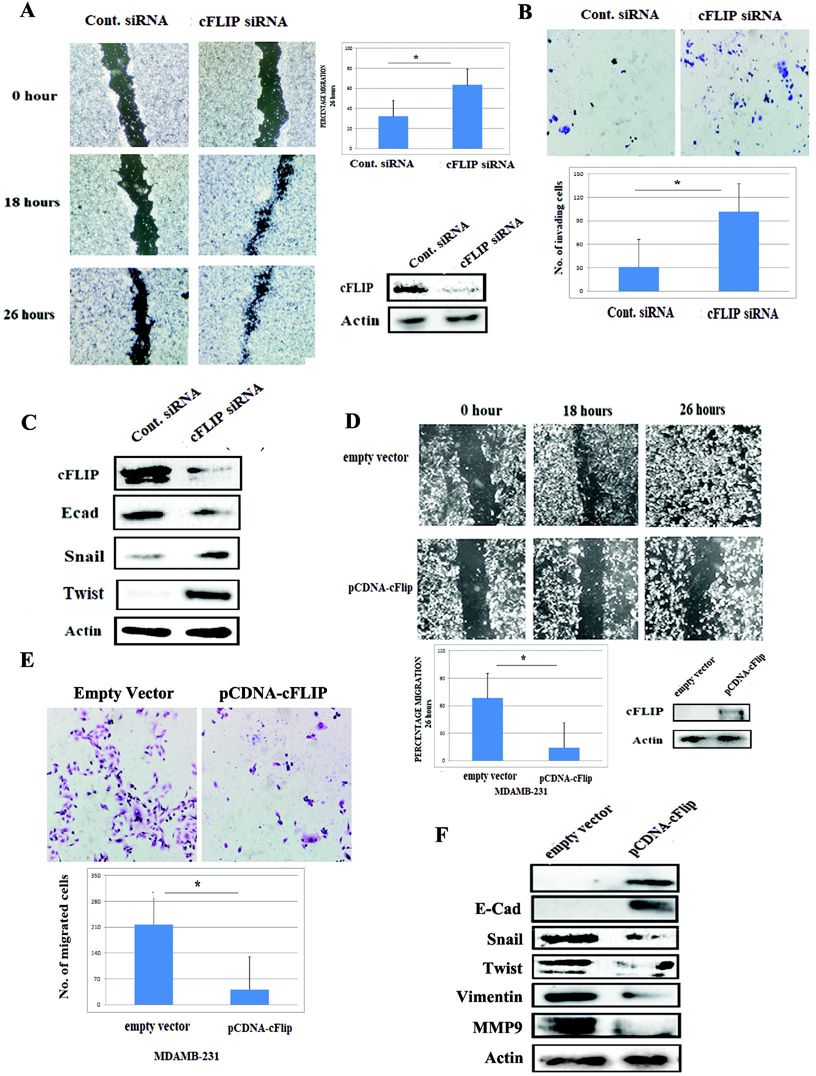
Involvement of cFLIP in migration and invasion properties of breast cancer cells. (A) Scratch assay was performed according to the standard protocol in MCF7 cells treated with control and cFLIP siRNA. Representative photographs of one random field is provided and the wound area was representeted in form of bar graph with mean ± SD. (B) Transwell assay was performed in the standard protocol in MCF7 cell lines with cFLIP knock down using siRNA and four random representative fields were photographed. Cells were counted and represented in form of bar graphs with mean ± SD. (C) The expression of indicated proteins were analysed using western blot from the control and cFLIP siRNA treated MCF7 cell lines. For loading control γ Actin was checked. (D) Scratch assay was performed according to the protocol in MDAMB-231 cells with cFLIP overexpression. Representative photographs of one random field is provided and the wound area was representeted in form of bar graph with mean ± SD. (E) Invasion assay was performed in the standard protocol in MDAMB231 cells with cFLIP overexpression and four random representative fields were photographed. Cells were counted and represented in form of bar graphs with mean ± SD. (F) The expression of indicated proteins were analysed using western blot from the control and cFLIP overexpressing MDAMB 231 cell lines. For loading control γ Actin was checked.

Since TNBC cell line MDA-MB-231 showed negligible intrinsic cFLIP expression, cFLIP was ectopically expressed to validate its anti-migratory/invasive characteristics. Therefore, MDA-MB-231 cells were transiently transfected with either empty pCDNA3 vector as control or pCDNA3-cFLIP plasmid and subjected to scratch assay and transwell migration assays. Scratch assay demonstrated that cells expressing vector control nearly completely filled the wound by 26 h, whereas cFLIP overexpression markedly hindered the cell migration (Fig. 3D). Next, transwell invasion assays with matrigel revealed that cFLIP overexpressed MDA-MB-231 cells had a marked 4 fold reduction in cell motility as compared to the cells transfected with vector control (Fig. 3E), further underlining the important role of cFLIP in the regulation of breast cancer cell migration. The ectopic expression of cFLIP was confirmed by western blot analysis using a part of these cells used in the above-mentioned experiments (Fig. 3E). In contrast to cFLIP knockdown in MCF7 cells, ectopic expression of cFLIP in TNBC line demonstrated significant downregulation of mesenchymal markers including Snail, Twist and Vimentin, while epithelial marker E-cad showed an opposite trend (Fig. 3F). Overall, these data provided compelling evidence of a negative regulatory role of cFLIP in metastasis and cell migration.

### cFLIP mediated autophagy inhibition plays a crucial role in modulating its anti-migratory ability of breast cancer cells

Since downregulation of cFLIP plays a pro-metastatic role in advanced stage breast cancer, we next investigated the antagonistic role of cFLIP in breast cancer metastasis. Accumulating evidence suggests paradoxical roles of cellular autophagy pathway in tumor-suppression as well as tumor-progression. Interestingly, cFLIP was shown to be associated with autophagy pathway (13,14). Therefore, we tried to identify whether cFLIP regulates its anti-metastatic potential of the breast cancer cells in an autophagy dependent manner. Since we demonstrated that the migration ability along with cFLIP expression distinctly differed in four breast cancer cell lines – including luminal line MCF7 along with three TNBC lines MDA-MB-231, MDA-MB-453 and MDA-MB-468, we further checked the autophagy markers in these cells by western blot analysis (Fig. 4A). Elevated lapidated LC3B-II and ATG12 expressions along with lower p62 level indicated enhanced autophagic flux in two TNBC lines MDA-MB-231 and MDA-MB-468 (Fig. 4A). In contrast, MCF7 cells with highest cFLIP expression and least migration ability exhibited negligible expression of autophagy markers. To further corroborate this finding, immunofluorescence analyses were performed with LC3B-II antibody in MCF7 and MDA-MB-231 cell lines (Fig. 4B). As similar to western blot analysis, immunofluorescence data also confirmed elevated LC3B-II expression in MDA-MB-231 cells as compared to MCF7 cells (Fig. 4B).

**Figure 4:**
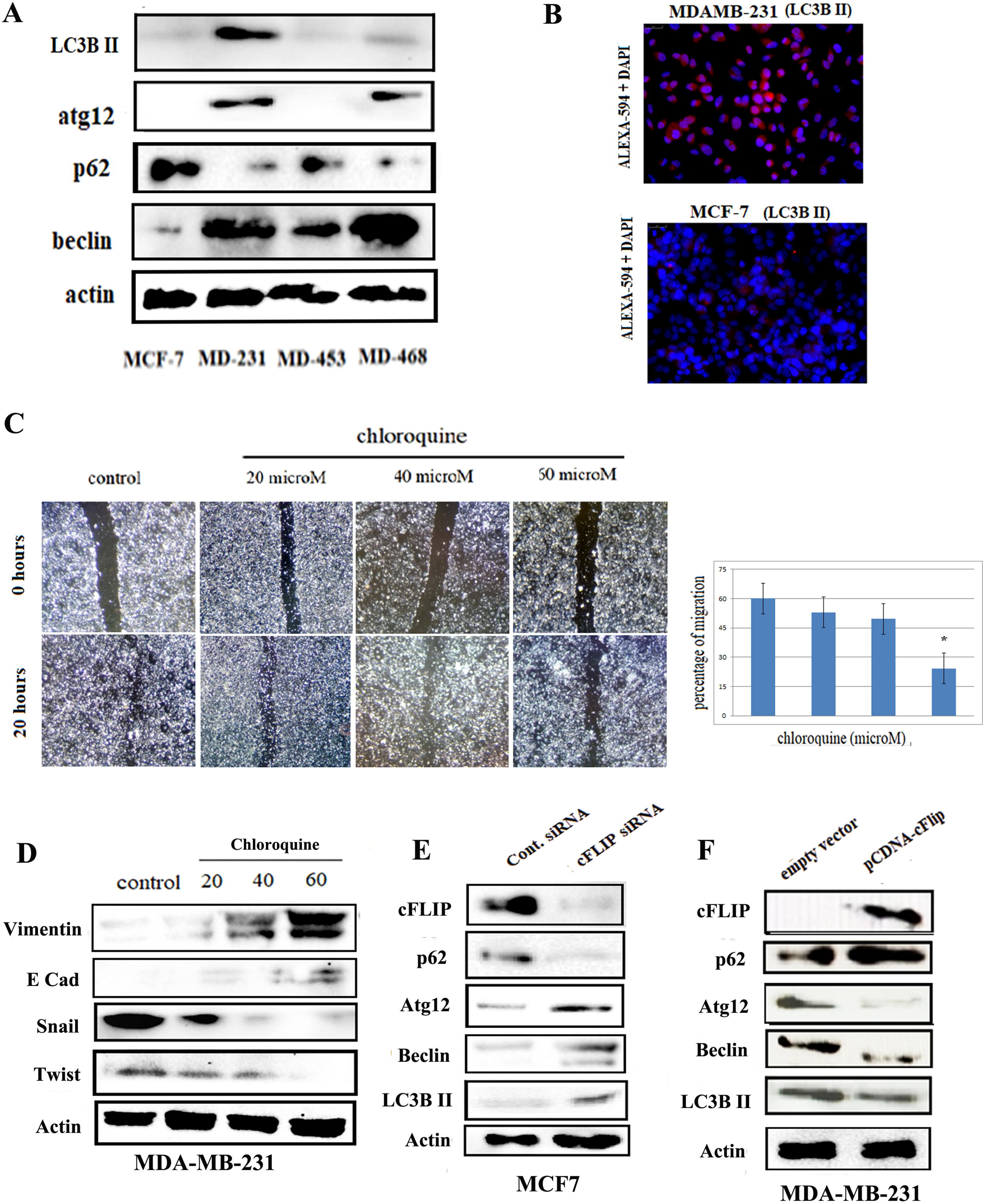
cFLIP having an autophagy mediated effect on modulating migratory ability of breast cancer cells. (A) The expressions of the autophagic markers were analysed using western blot technique from the four mentioned cell lines. For loading control γ Actin was checked. (B) MDAMB 231 and MCF7 cells were seeded for immunofluresence overnight, and fixed using 10% PFA onto the slides and incubated with anti LC3b II antibody overnight and stained with DAPI (Blue) and Alexa Fluro 594 (Red). (C) Scratch assay was performed according to the protocol in MDAMB 231 cells with control and indicated chloroquine concentration treatment. Representative photographs of one random field is provided and the wound area was representeted in form of bar graph with mean ± SD. (D) The expressions of the epithelial to mesenchymal transition markers were analysed using western blot technique from control and indicated dose of chloroquine treated MDAMB231 cells. For loading control γ Actin was checked. autonomous repeats. (E) The expressions of the autophagic markers were analysed using western blot technique from the MDAMB231 cells with indicated treatments. For loading control γ Actin was checked. (F) The expressions of the autophagic markers were analysed using western blot technique from the MCF7 cells with indicated treatments. For loading control γ Actin was checked.

In order to validate whether enhanced autophagy in MDA-MB-231 is critical for increased metastatic ability, wound healing/scratch assay was performed in the presence of increasing concentrations of chloroquine (Fig. 4C). Chloroquine is a well-established inhibitor of autophagy, which blocks autophagosome and lysosome fusion and slows down lysosomal acidification. The wound healing assay revealed that migration ability of cell was significantly reduced after chloroquine treatment compared with the DMSO control (Fig. 4C). In addition, chloroquine treatment resulted in increased epithelial marker E-cadherin and decreased mesenchymal markers Snail, Twist and Vimentin expression levels in a dose dependent manner (Fig. 4D), signifying that a functional autophagy plays a positive regulation in epithelial-to-mesenchymal transition and metastasis.

To further corroborate cFLIP mediated autophagy inhibition, MCF7 cells were knockdown for cFLIP using specific siRNA transfection, while MDA-MB-231 cells were transiently transfected with cFLIP expressing construct and subjected western blot analysis for autophagy markers (Fig. 4E and 4F, respectively). As compared to the control, cFLIP knockdown in MCF7 cells resulted in enhanced ATG12, Beclin 1 and lipidated LC3B-II expressions along with decreased p62 level (Fig. 4E). In contrast, ectopic expression of cFLIP in MDA-MB-231 cell line exhibited an opposite phenomenon (Fig. 4F). Taken together, our data indicated that cFLIP mediated autophagy inhibition plays an adverse impact on metastatic potential of breast cancer cells, which could be further exploited as a therapeutic strategy.

### p38 plays a role in cFLIP down regulation in triple negative breast cancer cell lines

Next, we investigated the possible underlying mechanistic pathway that regulates cFLIP down regulation in TNBC lines with elevated metastatic potential, particularly in MDA-MB-231 cells. Previously we demonstrated the involvement of MAP kinase pathways in the regulation of cFLIP expression in K562 CML cells (4). Moreover, studies suggested that cFLIP exerts its anti-apoptotic activity, at least in part, by inhibiting p38 MAPK activation. We therefore first checked the expression prolife of MAP kinase pathway members in the four breast cancer lines (Fig. 5A). The pattern of the pERK, pJNK did not show any correlation with that of cFLIP expression across the cell lines (Fig 5A). In contrast, the phosphorylation pattern of p38 clearly showed an inverse correlation with cFLIP expression pattern in these cell lines (Fig. 5A). The phosphorylation patterns of both p38 and its one of the downstream targets c-Jun, were significantly elevated in MDA-MB-231 cells with least cFLIP expression (Fig. 5A).

**Figure 5:**
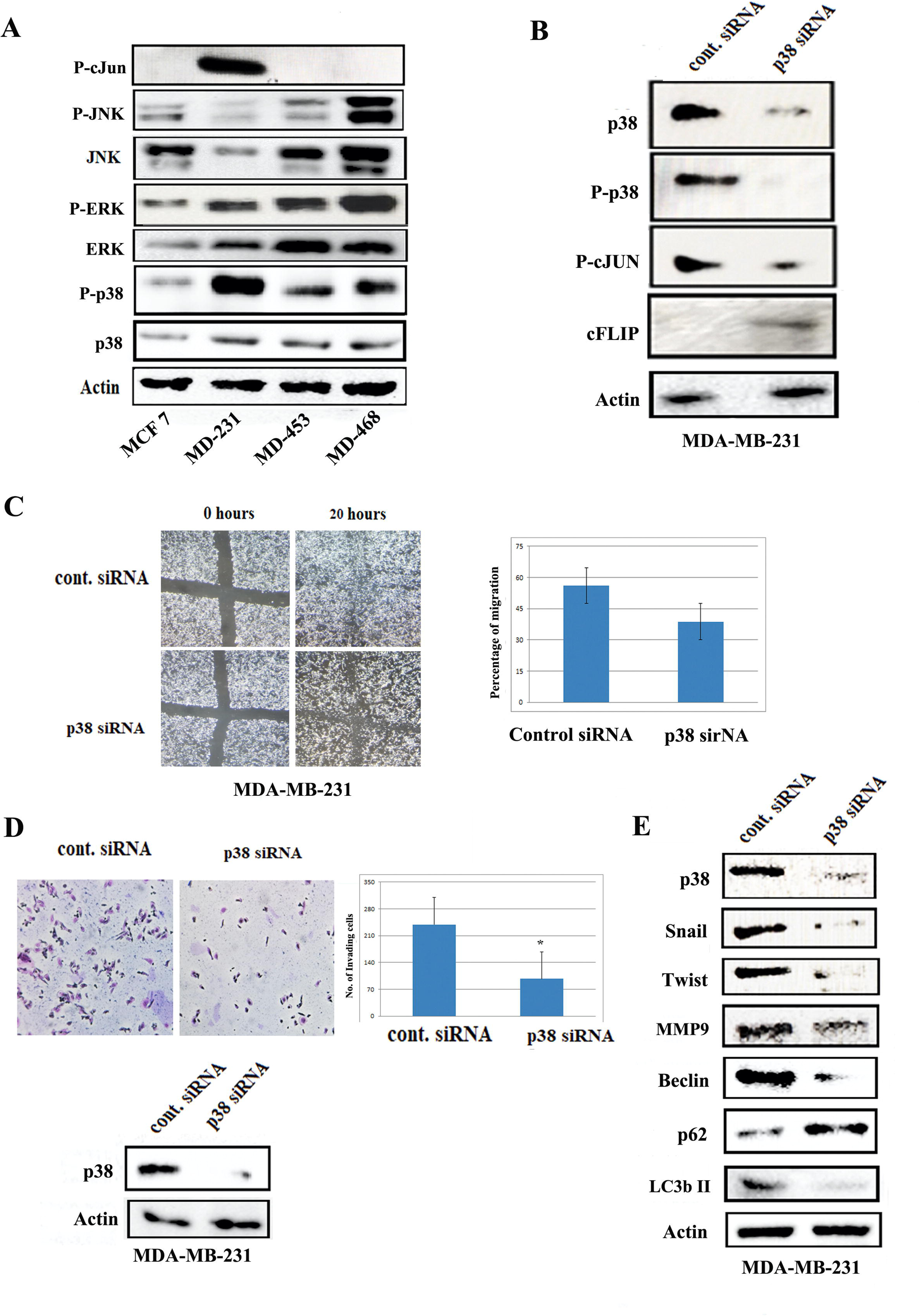
p38 mediated cFLIP downregulation in triple negetive breast cancer cell lines. (A) The expressions of the MAP kinase pathway proteins were analysed using western blot analysis from the four mentioned cell lines. For loading control γ Actin was checked. (B) The expression of mentioned proteins was analysed using western blot analysis from siRNA p38 knock-down MDAMB 231 cells. For loading control γ Actin was checked. (C) Scratch assay was performed according to the protocol in MDAMB 231 cells with control and p38 si RNA treatment. Representative photographs of one random field is provided and the wound area was representeted in form of bar graph with mean ± SD. Representative western blot confirms the knock down effect of the siRNA. (D) Transwell assay was performed in the standard protocol in MDAMB231 cells with siRNA mediated p38 knock down and four random representative fields were photographed. Cells were counted and represented in form of bar graphs with mean ± SD. Representative western blot confirms the knock down effect of the siRNA.

Inverse correlation of cFLIP and p-p38 expression prompted us to hypothesize that p38 may regulate cFLIP expression in breast cancer cells. In order to determine the possible involvement of p38 mediated cFLIP expression control, p38 was knockdown using specific siRNA in MDA-MB-231 cells and subjected to western blot analysis. In support of our hypothesis, a partial rescue of cFLIP expression was demonstrated in p38 knockdown MDA-MB-231 cells as compared to the control line (Fig. 5B). A marked reduction of phosphorylated c-Jun in p38 knockdown cells further authenticated functional knockdown of p38 MAP Kinase pathway (Fig. 5B). To confirm the role of p38 in regulating cFLIP anti-metastatic potential, wound healing/scratch and transwell-invasion assay were performed using MDA-MB-231 cell lines transfected with either scramble siRNA or p38 specific siRNAs (Fig. 5C and 5D, respectively). Inhibition of p38 pathway resulted in delayed wound healing and cell-invasion as compared to the control line (Fig. 5C, 5D). In addition, p38 knockdown in MDA-MB-231 cell line resulted in stark decreased of mesenchymal markers – Snail, Twist and MMP9 (Fig. 5E). Since cFLIP negatively regulates autophagy, we also evaluated autophagy markers in p38 knockdown cells. The results demonstrated that p38 knockdown caused a significant reduction of Beclin and LC3B-II levels, whereas p62 level was increased, indicating inhibition of autophagy flux(Fig. 5E). Altogether, these results indicated that p38 activation plays a significant role in reduced cFLIP expression and increased autophagy, which in turn promotes metastatic ability in TNBC.

### IL6 promotes the migratory ability of breast cancer cells by inhibiting cFLIP expression

So far, we demonstrated that advanced stage breast cancer, TNBC had increased metastatic potential via activation of p38 MAP kinase pathway and suppression of cFLIP expression as compared to non-aggressive luminal breast cancer. A large body of evidence suggests that concentration of myeloid-derived suppressor cells (MDSCs) and tumor-associated macrophages (TAMs) in the cancer microenvironment promotes tumor growth either directly by augmenting tumor cell proliferation and survival or indirectly by building an immunosuppressive microenvironment (15,16). IL6 is one of the most prominent cytokines secreted by these two cells (17). We hypothesised that IL6 might play a role in p38 activation, cFLIP down regulation and subsequent events which lead to increased metastatic potential. Moreover, it is not known whether there is any correlation between IL6 and cFLIP.

To prove our hypothesis, we first checked the endogenous expression of IL6 receptor (IL6R) among the four breast cancer lines utilized in previous studies since IL6 exerts its biological functions through its receptor IL6R and the later would be increased in a positive feedback manner with increased availability of IL6 and subsequent binding to its receptor. Both PCR and western blot analyses demonstrated a similar pattern of IL6R expression that was observed for p38 in these cell lines (Figs. 6A and 6B, respectively), indicating that IL6 might be involved in regulating aggressiveness of TNBC lines MDA-MB-231 and MDA-MB-468 with negligible cFLIP expression as compared to the other two lines MCF7 and MDA-MB-453. The inverse correlation of IL6R and cFLIP in different breast cancer cell lines prompted us to further investigate the potential role of IL6 in cFLIP suppression. To this end, MCF7 cells were treated with 20 pg/ml recombinant IL6 and subjected to western blot analysis (Fig. 6B). IL6 treatment resulted in a sharp decrease in cFLIP and E-cad levels (Fig. 6B), indicating that IL6 mediated metastatic promotion of MCF7 cells through downregulation of cFLIP expression.

**Figure 6:**
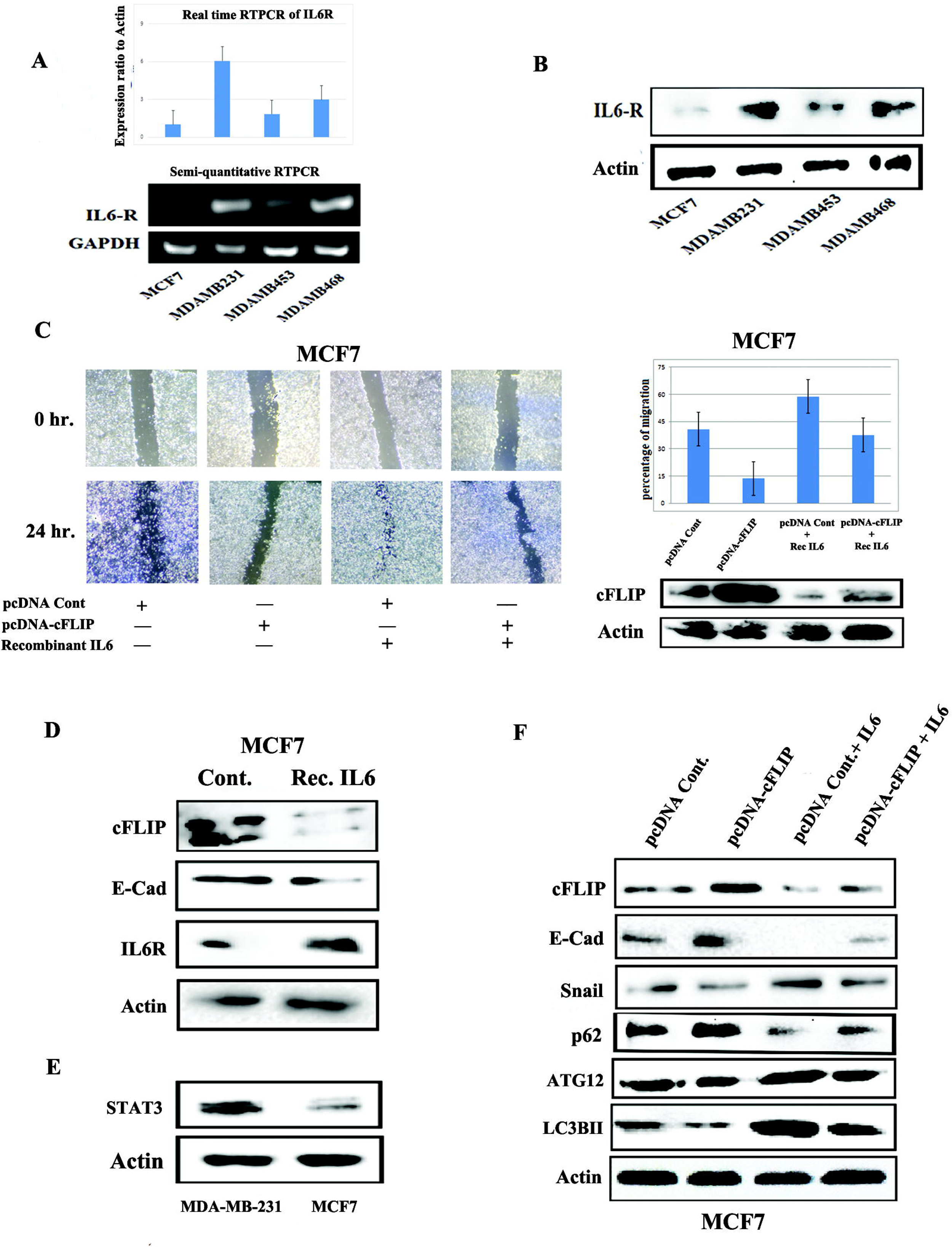
Il6 modulates the migratory ability of breast cancer cells by influencing the cFLIP expression. (A) Semi quantitative PCR was performed for 30 cycles and Realtime PCR for 40 cycles were performed to check the expression of IL6R mRNA in the four mentioned cell lines. For loading control γ Actin and GAPDH was checked. Densitometric analysis data was represented graphically normalised with actin and indicated ± SD. (B) The expressions of indicated protein was checked using western blot technique from control and treated MCF7 cells. For loading control γ Actin was checked. (C) Scratch assay was performed according to the protocol in MCF7 cells with indicated treatments. Representative photographs of one random field is provided and the wound area was representeted in form of bar graph with mean ± SD. Representative western blot confirms the knock down effect of the siRNA. (D) The expression of mentioned proteins was analysed using western blot analysis from the MCF7 cells with the indicated treatments. For loading control γ Actin was checked. (E) The endogenous expression of mentioned protein was analysed using western blot analysis from the MCF7 and MDAMB231 cells. For loading control γ Actin was checked. (F) The expressions of indicated protein was checked using western blot technique from control and mentioned treatments in MCF7 cells. For loading control γ Actin was checked.

To further corroborate IL6-cFLIP axis in breast cancer metastasis, scratch assay was performed using MCF7 cells transiently transfected with either vector control or cFLIP expressing construct followed by IL6 treatment (Fig. 6C). The results revealed that the IL6 treatment caused significant increment of cell migration in MCF7 cells transfected with empty plasmid as compared to cFLIP overexpressing MCF7 cells, indicating that cFLIP overexpression might impede IL6 mediated migratory ability of breast cancer cells. A parallel experimental set up was also conducted and subjected to western blot analysis for checking expression pattern of both autophagy and EMT markers (Fig. 6D). The data clearly showed that IL6 treatment resulted in augmentation of autophagy flux as indicated by increased LC3B-II and ATG12 expressions and decreased p62 level in MCF7 cells with vector control as compared to cFLIP overexpressing MCF cells (Fig. 6D). Likewise, expression pattern of metastasis markers – Snail and E-cad further supported the findings from scratch assay (Fig. 6D). In sum, these findings suggested that IL6 promotes breast cancer invasiveness through down regulating cFLIP expression. The expression of STAT3 was also checked in the two cell lines namely luminal less aggressive MCF7 and more aggressive triple negative MDAMB 231 (Fig. 6E) and an elevated level of STAT3 was found in MDA-MB-231 cell. STAT3 is a bonafide downstream candidate for IL6R signalling. Therefore, it can be concluded that MDAMB231 shows inherently more IL6 mediated signalling capability. Now we hypothesised that does IL6 mediates its’ effect on metastasis through cFLIP and autophagy. To put light in that context we checked the expression of the autophagy markers in the presence of recombinant human IL6 in cFLIP overexpression in the MCF7 cell line (Fig. 6F). And the expression patterns of the autophagy and EMT markers showed that the IL6 mediated effect on autophagy and metastasis was reversed to some extent on overexpression of cFLIP in the MCF7 cell line.

Thus, here we have shown that IL6 downregulate the expression of cFLIP through modulating the p38 activation and cFLIP downrehulation is seen to play an important role on upregulation of autophagy and increase in the metastatic ability of the late stage breast cancer cells (Fig 7).

**Figure 7:**
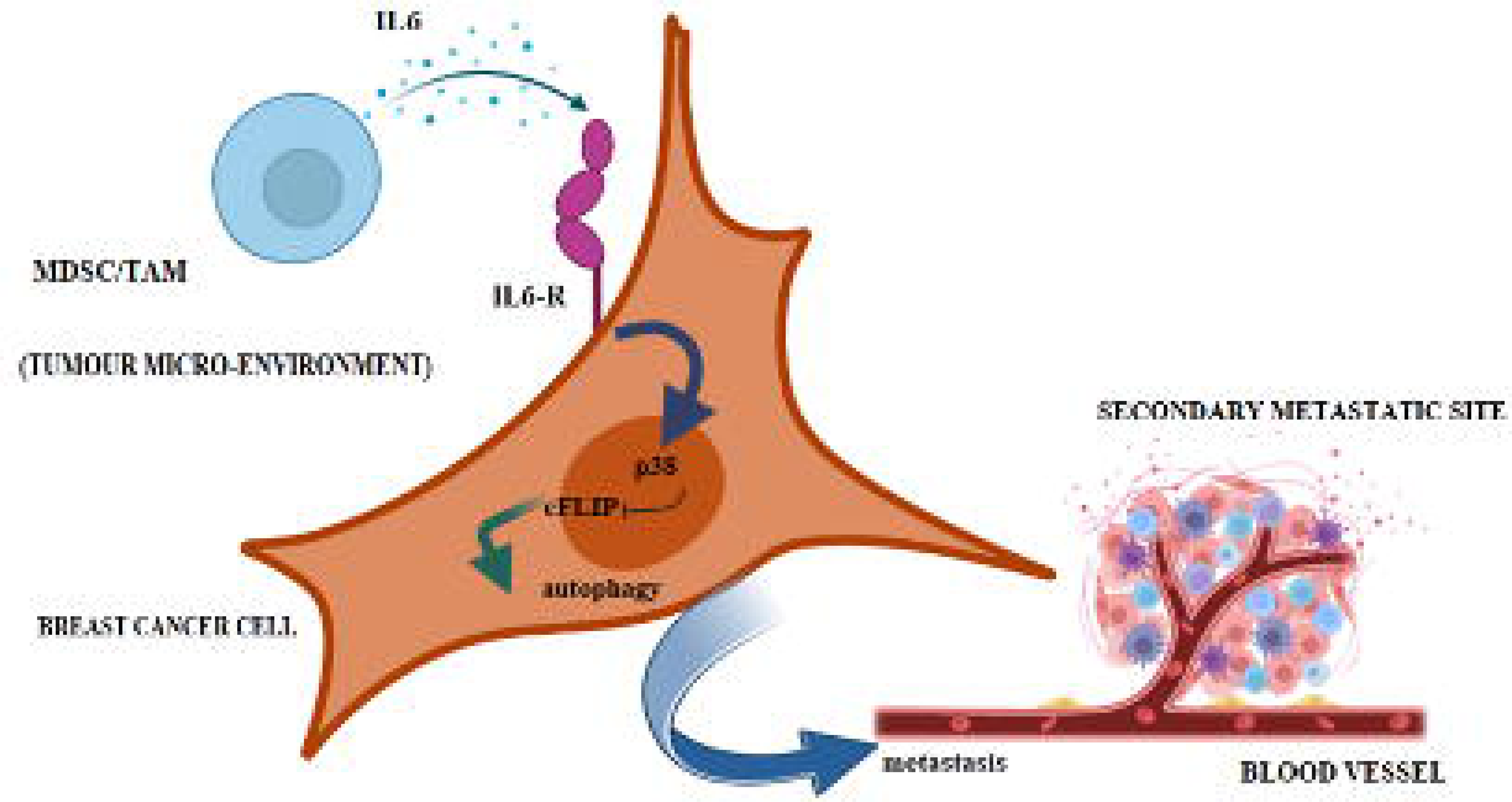
Mechanisms of IL6 mediated regulation of metastasis by downregulation of cFLIP. IL6, one of the prominent cytokines secreted by tumour infiltrating immune cells as well as by tumor associated fibroblasts activates the p38 pathway. P38 downregulates the expression of cFLIP and thus enhances autophagy. Increased autophagy enhances the invasion and migration capacity of aggressive breast cancer cells.

## Discussion

cFLIP (cellular FLICE-like inhibitory protein) has regulatory functions in apoptosis, necroptosis, autophagy, and inflammation (17). cFLIP is also involved in innate and adaptive immunity as well as during embryonic development (18). Up regulation of cFLIP has been reported to contribute to drug resistance and tumour progression in many types of cancers, including prostate, colorectal, bladder urothelial, cervical and lymphomas (19). We have recently shown that cFLIP regulation by hydroxychavicol makes imatinib-resistant CML cells sensitive to TRAIL (20). In addition, we have also shown that cFLIP is regulated via ROS in a proteasomal degradation pathway in imatinib-resistant cells. In another work, we observed that cFLIP is expressed more in imatinib resistant K562 CML cells as compared to sensitive cells (unpublished data). In an aim to gain knowledge of cFLIP expression status in different stages of breast cancers, TCGA analyses revealed lower cFLIP expression in metastatic stages of breast cancer. Given the association of lower levels of cFLIP during metastasis, we hypothesized that low levels of cFLIP may be associated with poor clinical outcomes. Indeed, analysis of data derived from breast cancer patients revealed that lower levels of cFLIP expression was linked to poor patient outcomes of both luminal and TNBC categories. *In vitro* experiments with different cell line models subsequently validated that cFLIP levels are significantly downregulated in more aggressive breast cancer types - TNBC cell lines namely MDA-MB-231 when compared to luminal types like MCF7.

This study deals with a Bonafede anti-apoptotic protein cFLIP. Being an antiapoptotic protein it’s upregulation would stop cells from apoptotic death which is necessary for cancer progression. Now based on these observations, checking the link between metastasis and cFLIP became the next important search. Therefore, the migrating and invasive ability of the different breast cancer cell lines were checked first. The TNBC cells namely MDA-MB-231 and MDA-MB-468 showed higher migratory and invasive ability than that of luminal MCF7 cells. The epithelial to mesenchymal transition markers supports their migratory potentials found in the scratch assay. Interestingly, MDA-MB 453, although known as metastatic cell line didn’t show much migration in the scratch assay. Moreover, it was seen that cFLIP expression levels are higher in the cells with less invasive and migratory ability i.e. MCF7 and MDA-MB-453 which further supported our hypothesis that cFLIP downregulation is somehow involved in the increase in invasive or metastatic ability of the breast cancer cells.

Next, to ascertain the effect of cFLIP on metastasis, the protein was knocked down in MCF7 cell line using siRNA and the migration rate was checked with scratch assay. MCF was used because this cell line has inherent high expression of the cFLIP protein. The expressions of the EMT markers were also checked in this knocked down condition. We found that the migratory and invasive ability of the cells was increased along with increase in mesenchymal markers and decrease in epithelial markers on decreasing the expression of cFLIP. On the other hand, cFLIP was ectopically overexpressed in MDA-MB-231 cells where the expression of cFLIP is inherently almost absent and the same wound healing assay was performed. It was seen that with overexpression of cFLIP the migratory and invasive ability of the cells decreased which also was reflected in epithelial mesenchymal markers expression pattern. Therefore, from these data it can be concluded that cFLIP has a role to play in the increase in metastasis of the breast cancer cells.

cFLIP being an anti-apoptotic protein is known to help cell survival. But here a noncanonical role of cFLIP was observed and consequently a question arises that how cFLIP can bring about the change in metastatic ability of the breast cancer cells. Autophagy, a highly conserved self-degradative process, and a key process in cellular stress responses has been explored previously and found to have functions in cancer metastasis (21). This is of specific interest given the scarcity of effective therapies for metastatic patients. Autophagy is induced in aggressive stages for various cancers and metastasis and along with evidences from genetically engineered mice models and other experimental metastatic models, roles of autophagy in almost every stages of metastasis has been found. (22). We have also found some critical link between cFLIP and autophagy in one of our studies (manuscript under preparation). So, in the context of this work firstly the status of autophagy was checked in the four breast cancer cell lines. It was found, as expected, that autophagy markers LC3IIb, Atg12 increased and p62 get reduced in MDA-MB-231 and MDA-MB-468 cell lines which signified that autophagy is upregulated there. Strikingly, the autophagy markers showed just the opposite trend of that of cFLIP, means, autophagy was upregulated in the cell lines with less cFLIP expression. Moreover, autophagy inhibitor chloroquine also decreased the metastasis in MDA-MB-231 cells which was observed by scratch assay. The EMT marker expressions pattern also corroborated our findings. Therefore, it can be concluded that autophagy upregulation has a positive role in the metastatic activity. This autophagy upregulation is directly connected to cFLIP downregulation because when we knocked down cFLIP in MCF7 cells, it increased autophagy which is evident by increased autophagic markers. On the other hand, when we ectopically expressed cFLIP the autophagy got downregulated in MDA-MB-231 cells. Therefore, it was confirmed that advanced stage cancer cells downregulate cFLIP and as a result of this downregulation autophagy get upregulated which in turn increase the metastatic ability of the cell.

The obvious questions then comes is how the expression of cFLIP is controlled in the breast cancer cell lines. Previous research from our lab has showed the involvement of MAPK pathway in controlling the expression of cFLIP, XIAP. Therefore, Bonafede MAPK pathway proteins were checked in the four breast cancer cell lines. The active forms, i.e, the phosphorylation form of ERK and JNK did not show any correlation with that of the expression of flip in the four cell lines. But interestingly the phospho-P38 showed just the reverse activation pattern of that of cFLIP in the cell lines, i.e phospho-p38 was high in the cell lines with less cFLIP expression. Moreover, one of the downstream transcription factors that get phosphorylated and activated by p38, i.e P-cJun, also showed highest activation in MDA-MB-231. Interestingly, JNK, P-JNK, Erk and P-Erk showed some differential expressions and activation patterns. However, we have not explored them further as the pattern of either the phosphorylation or expression didn’t correlate with that of expression patterns of cFLIP in these four cell lines. Next, when we knocked down p38 by siRNA in the MDA-MB-231 cell line, cFLIP level increased and cell migration and invasion decreased as observed by scratch assay and transwell invasion assays respectively. Not only that the p38 knockdown also increased mesenchymal markers and decreased epithelial markers to confirm the increase metastasis along with increase in autophagy.

The effects of the cytokines present in the tumour microenvironment (TME) in tumour progression are well documented (23). Among many of such cytokines, IL6 is an eminent member which is known to modulate several pathways in the different cancer cells which cause tumour progression (24). TNBC is the most challenged one to treat because of its invasiveness and increasing chemo resistance. Il6 itself sometimes play important roles in resistance of TNBC which make it harder to target (25). IL6 is well known to modulate the JAK/STAT, ERK and Akt pathway mainly to manipulate the cells (26). But there are some cell specific contexts where involvement of other pathways, are also evident (27). As mentioned before, several cells like TAM and MDSC are present inside the tumour microenvironment which augment the tumour progression and make a fool of the immune system so that they are unable to spot the cancer cells (28,29). The mode of action of these pro-tumorigenic cells includes the secretion of cytokines that help tumour cells to grow (30). These two cells and some of these pro-tumorigenic cytokines, increase in advanced stages of cancer (31). One of the prominent cytokine inside tumor microenvironment is IL6 which is secreted by Tumor associated macrophage, Tumor associated fibroblast, by the tumour cells themselves, and by stromal cells (32). Moreover, Shen eta al showed that Interleukin-6 induces Akt and p38 MAPK activation and migration of fibroblast in non-diabetic but not in diabetic mice (33). Therefore, it was worth to check this cytokine here to search whether it has any role in cFLIP downregulation and increase in metastasis as both of these are late-stage events. Il6 mediates its effect through Il6R, Il6 receptor. Il6R can be soluble or membrane bound (34). Sometimes a feed-forward loop is also observed in this pathway that is binding of ligand to the receptor increases the expression of the receptors and further signalling (35). It was found that the IL6R expression was high at RNA level in the TNBC lines with respect to that of the luminal type. Moreover, STAT3, the downstream target of IL6, was also found high in MDA-MB-231. Interestingly, this corroborates previous finding that stress activated protein kinase p38 plays a role in IL-6 induced activation of transcription of STAT3 (36). Recombinant IL6 treatment showed a decrease in the cFLIP level in the MCF7 cell line where cFLIP level is endogenously high. Now, the role of IL6 in metastasis was confirmed when we found that IL6 treatment in MCF7 cells increased metastasis and the later was suppressed when cFLIP was overexpressed. The EMT and the autophagic markers were checked in the same experimental condition and it was found that the IL6 increased mesenchymal markers and autophagy in cFLIP dependent manner. Not only that, our p38 knockdown data perfectly concluded that IL6 mediated cFLIP downregulation is p38 dependent.

Therefore, in conclusion, this work explained an axis which starts from pro-tumorigenic cytokine IL6 that are abundant in the tumour microenvironment, and ultimately ends in the increase in metastatic potentials of breast cancer cells. IL6 activated the p38 pathway in the advanced breast cancer stages to decrease the expression of cFLIP that is ultimately bringing an alteration of the autophagic flux and EMT markers in the cells and resulting in their more aggressive nature. Thus, this study, for the first time, shows a non-canonical pro-metstatic role of a Bonafede antiapoptotic protein cFLIP. Moreover, it also uncovers an IL6, p38 mediated axis that partially is responsible for maintaining the expression of the cFLIP protein which was not identified yet.

## Acknowledgement

We would like to sincerely acknowledge Prof. Mikhiko Naito, National Institute of Health Sciences, Tokyo, Japan for providing pcDNA3-cFLIP. This work was financially supported by the SERB, Department of Science and Technology, Govt. of India (EEQ12019/000153); ICMR, Govt of India (2021–11421) and DBT BUILDER (BT/INF/22/SP45088/2022).

## Conflict of Interest

The Authors declare no conflict of interest.

## Notes

### Competing Interest Statement

The authors have declared no competing interest.

### Summary of Updates

The figures and manuscript, specially the introduction part and some missed refernces ahve been added

